# Various Cell Types in the Bone Marrow sustain Primary B-Cell Precursor Acute Lymphoblastic Leukemia

**DOI:** 10.1101/2023.05.25.542080

**Authors:** Mandy W. E. Smeets, Cesca van de Ven, Jan Orsel, Caitlin E. J. Reichert, Aurélie Boeree, Judith Boer, Monique L. den Boer

## Abstract

B-cell precursor acute lymphoblastic leukemia (BCP-ALL) originates from the bone marrow, which besides hematopoietic cells also contains different (supportive) cell types, including mesenchymal stromal cells (MSCs), osteocytes, chondrocytes, fibroblasts, and adipocytes. These (supportive) cell types create a bone marrow microenvironment that facilitates leukemogenesis and provide a survival benefit to leukemic cells that also affects the response to chemotherapeutic drugs. We here show that the survival benefit provided by supportive tissues does not depend on the type of supportive cells and does not differ between supportive cells collected at the time of full leukemia (diagnosis), at end of consolidation therapy or from healthy controls. The need for supportive cells, however, clearly differed between subtypes of BCP-ALL, with *BCR-ABL1*-positive and *TCF3-PBX1*-positive subtypes being the most dependent on this support. Various supportive cell types provided a survival benefit to BCP-ALL cells, with a median benefit of 18% (chondrocytes) and 30-36% (MSCs, osteocytes, and fibroblasts), which was not observed for mature adipocytes. This benefit was direct cell-cell contact dependent and decreased upon physical separation of cell populations in a transwell setting. BCP-ALL cells in contrast to MSCs and the other supportive tissue types hardly produce cyto-/chemokines. The secretome of MSCs changed upon co-culture with BCP-ALL cells resulting in 1.2-fold to 1.4-fold (median) higher levels for the cyto-/chemokines IL6, CCL22, CXCL10, and CXCL5. Together, these data suggest that BCP-ALL cells manipulate different components of the bone marrow supportive tissues via direct cell-cell contact which favors the survival of leukemic cells. This strengthens our earlier observation that BCP-ALL cells hijack the bone marrow microenvironment and offers the perspective that interference with this stromal interaction and/or released cyto-/chemokines may be of additive value in the treatment of BCP-ALL.

## Introduction

Acute lymphoblastic leukemia (ALL) is the most common childhood malignancy ^1,2^. The majority of pediatric cases suffer from B-cell precursor (BCP) ALL (85%) which is characterized by recurrent genetic aberrations affecting B-cell transcription factors and tyrosine kinase genes ^3,4^. BCP-ALL can be divided into favorable cytogenetic subtypes, e.g., *ETV6-RUNX1*, high hyperdiploidy (51-67 chromosomes) and *TCF3-PBX1*, and unfavorable subtypes including ABL-class/*BCR-ABL1* and *KMT2A*-rearranged cases ^5–7^. Cases without these established genetic abnormalities are called “B-other” in this paper, and represent a highly heterogeneous group of patients with various types of lesions and variable outcome ^4^. Although the introduction of risk-adjusted treatment protocols based on cytogenetic subtype and response to initial therapy (minimal residual disease, MRD) has improved the outcome of pediatric ALL to a 5-year event-free survival rate of 90% ^5,8,9^, BCP-ALL still represents a main cause of cancer-related mortality in children ^9,10^.

For many years, research was focused on leukemic cells ^11^. However, in recent years the emphasis shifts more and more towards leukemic cells and the bone marrow microenvironment that encompasses these malignant cells ^12–15^. This leukemic microenvironment contains different cell types including other, non-leukemic immune cells, blood vessels/endothelial cells and supportive cell types like osteoblasts, fibroblasts and mesenchymal stromal cells (MSCs) ^11,16^. The latter are fibroblast-like, multipotent cells that can differentiate into different mesodermal lineages, including adipocytes, osteocytes, and chondrocytes ^17,18^. MSCs constitute a major component of the microenvironment in the bone marrow ^19^. We previously showed that these MSCs support the viability of primary BCP-ALL cells and contribute to resistance of leukemic cells to prednisolone, the active form of the major chemotherapeutic drug prednisone used in the treatment of BCP-ALL ^15,20^. Disruption of this interaction reduced the viability and sensitized BCP-ALL cells to drugs ^15^. In the present study, we investigated which other cell types of the bone marrow microenvironment also support BCP-ALL cells and whether this support differs between cytogenetic subtypes of BCP-ALL and/or in the type of cyto-/chemokines being secreted by these cell types.

## Materials and Methods

### Mesenchymal stromal cells

Bone marrow-derived MSCs were obtained from pediatric bone marrow aspirates at diagnosis (day 0), after consolidation treatment (day 79), at leukemic relapse and from normal, healthy condition. MSCs were selected based on adherence capacity, followed by confirmation of negativity for surface markers CD19/CD34/CD45, and positivity for CD44/CD54/STRO-1/CD73/CD90/CD105/CD146/CD166. MSCs were passaged twice a week in DMEM low glucose(1g/L)/pyruvate/HEPES medium (Gibco, ThermoFisher Scientific, Bleiswijk, The Netherlands) supplemented with 15% fetal calf serum (FCS) (Bodinco BV, Alkmaar, The Netherlands), amphotericin B (1:166; 250μg/ml, Gibco), gentamicin (1:1000; 50ng/ml, Gibco), fresh 1ng/ml recombinant human fibroblast growth factor (FGF) basic (BioRad, Veenendaal, The Netherlands) and 0.1mM L-ascorbic acid (Sigma Aldrich, Zwijndrecht, The Netherlands). MSCs were cultured at 37°C/5%CO_2_ and used for experiments until passage 10. MSCs from 12 different (ALL or healthy) donors were used (***Supplementary table 1***).

### Fibroblasts

Human Dermal Fibroblasts (HDFα) were purchased at Gibco, ThermoFisher Scientific (#C0135C). Cells were passaged twice a week and were maintained in DMEM low glucose(1g/L)/pyruvate/HEPES medium (Gibco) supplemented with 10% FCS (Bodinco), amphotericin B (1:166; 250μg/ml, Gibco), gentamicin (1:1000; 50ng/ml, Gibco). Fibroblasts were cultured at 37°C/5%CO_2_ and used for experiments until passage 35.

### Primary BCP-ALL cells

Primary BCP-ALL cells were collected from bone marrow samples originating from children with newly diagnosed BCP-ALL (1-18 years). This study was performed in agreement with the Institutional Review Board, and written informed consent was given by patients, parents and/or guardians. Isolation and processing of the leukemic blasts was performed by density gradient centrifugation using Lymphoprep (Nycomed Pharma, Oslo, Norway) for 15 minutes at 1500rpm ^21^. Normal hematopoietic cells were depleted using magnetic beads coupled to monoclonal antibodies to enrich for leukemic cells, resulting in ≥90% blasts prior to experiments. Primary BCP-ALL cells were kept in (short-term) culture at 37°C/5%CO_2_ using RPMI-1640 Dutch Modified medium (Gibco) containing 20% FCS, 2% PSF (penicillin 5.000U/ml, streptomycin 5.000U/ml, fungizone 250μg/ml, Gibco, ThermoFisher Scientific), gentamicin (0.2mg/ml, Gibco), 1% insulin-transferrin-selenium (ITS) (Sigma Aldrich), and 2mM L-glutamine (Life Technologies, ThermoFisher Scientific). A total of 42 unique primary BCP-ALL samples was used for this study, including 14 *ETV6-RUNX1*, 5 high hyperdiploid, 7 *TCF3-PBX1*, 9 B-other, 5 *BCR-ABL1*, and 2 *KMT2A*-rearranged BCP-ALL cases (***Supplementary table 2***).

### MSC differentiation assay

MSCs were seeded in 24-well plates at a density of 21.000 cells/cm^2^ for differentiation into adipocytes and 5.000 cells/cm^2^ for differentiation into osteocytes or chondrocytes. For differentiation staining, chondrocytes were seeded in parallel on 12mm coverslips (Electron Microscopy Sciences, Hatfield, PA, USA). Twenty-four hours after seeding MSCs, DMEM medium was replaced with adipogenic, osteogenic or chondrogenic induction medium. Control wells received DMEM medium without any adipo-, osteo-, or chondrogenic stimuli. Adipogenic medium consisted of DMEM high glucose(4.5g/L)/pyruvate/HEPES medium (Gibco) supplemented with 10% FCS (Bodinco), amphotericin B (1:166; 250μg/ml, Gibco), gentamicin (1:1000; 50ng/ml, Gibco), and freshly added indomethacin (0.2mM, Sigma Aldrich), insulin (1:1000; 11mg/ml, Sigma Aldrich), 3-isobutyl-1-methylxanthine (IBMX, 0.5mM, Sigma Aldrich), and dexamethasone (1μM, kindly provided by the Erasmus MC, Rotterdam, The Netherlands). Osteogenic medium consisted of DMEM low glucose(1g/L)/pyruvate/HEPES medium (Gibco) supplemented with 10% FCS (Bodinco), amphotericin B (1:166; 250μg/ml, Gibco), gentamicin (1:1000; 50ng/ml, Gibco), and freshly added β-glycerolphosphate (10mM, Sigma Aldrich), ascorbic acid (50 μM, Sigma Aldrich) and dexamethasone (1μM, Erasmus MC). Chondrogenic medium consisted of DMEM high glucose (4.5g/L)/ pyruvate/HEPES medium (Gibco) without FCS supplemented with amphotericin B (1:166; 250μg/ml, Gibco), gentamicin (1:1000; 50ng/ml, Gibco), and freshly added L-ascorbic acid (50 μM, Sigma Aldrich), L-Proline (0.35mM, Sigma Aldrich), human TGFβ3 (0.5mM, PreproTech, Rocky Hill, NJ, USA), and dexamethasone (1μM, Erasmus MC). Cells were cultured at 37°C/5%CO_2_, and medium was replaced twice a week for 21 days.

### Histochemistry of differentiated cells

The differentiation of patients’ derived MSCs into adipocytes, osteocytes and chondrocytes was confirmed by microscopic evaluation of histochemistry stained slides. For adipocytes, cells were fixed with 10% formalin (Sigma Aldrich) and stained with freshly made Oil Red O solution (2 parts Milli-Q and 3 parts Oil Red O, Sigma Aldrich). Osteocytes were fixed with ice-cold 70% ethanol at 4°C and stained with freshly prepared supernatant of saturated Alizarin Red S (Sigma Aldrich, 5mg/ml in Milli-Q). Images from adipo- and osteocytes were captured using a DMi1 microscope (Leica, Wetzlar, Germany). For chondrocytes, cells on the coverslips were fixed with 4% paraformaldehyde (Merck Millipore, Amsterdam, the Netherlands). Cells were permeabilized with 0.1% Triton-X100 (Sigma Aldrich), blocked with confocal laser scanning microscopy (CLSM; 1xPBS/3% bovine serum albumin/10 mM glycine) buffer and incubated overnight with Collagen II monoclonal antibody (2B1.5) (1:200, Invitrogen, ThermoFisher Scientific) at 4°C. Coverslips were incubated with secondary goat-anti-mouse AF647 antibody (1:400, Cell Signaling Technology, Danvers, MA, USA) and anti-phalloidin-AF488 (1:1000, ThermoFisher Scientific). Coverslips were placed on a microscopy slide containing 3μl ProLong Glass Antifade Mountant with NucBlue Stain (ThermoFisher Scientific). Imaging of chondrocytes was performed with a DM6 microscope (Leica) and analyzed by using ImageJ (version 2.0.0). MSCs differentiated efficiently until passage 8 for all lineages, while gradual loss of differentiation potential was evident from passage 12 (***Supplementary table 3***). MSCs were therefore used until passage 10 for subsequent (co-culture) experiments.

### Co-culture of primary BCP-ALL cells with MSCs

Primary BCP-ALL cells (1×10^6^ cells/ml) were co-cultured with primary MSCs (5×10^4^) for 5 days in a 24-well plate at 37°C/5%CO_2_. The percentage of viable leukemic cells was determined by staining with Brilliant Violet 421 anti-human CD19 (Biolegend, San Diego, CA, USA), FITC Annexin V (Biolegend), and propidium iodide (PI; Sigma Aldrich) as previously described ^15^, by flow cytometry using a MACSQuant analyzer (Miltenyi Biotec, Bergisch Gladbach, Germany). Viability of ALL cells in mono-culture was determined on day 0 (after thawing cells) and day 5 (after culture). Co-culture benefit (day 5) was determined by calculating the percentage of viable ALL cells in co-culture with MSCs minus the percentage of viable ALL cells cultured in parallel as mono-culture (= percent point; pp). Supernatant was collected after 40 hours of mono- and ALL-MSC co-culture and centrifuged for 10 minutes at 1200rpm to remove cellular debris. Supernatant was immediately stored at -80°C until quantification of cyto-/chemokine levels by Luminex technology (see below).

### Co-culture of primary BCP-ALL and different types of supportive cells

MSCs or fibroblasts (3.5×10^4^) were seeded in 24-well plates in DMEM medium (15 or 10% FCS, respectively), and cultured at 37°C/5%CO_2_. After 24 hours, 700μl fresh medium was added to the MSCs, fibroblasts and 21-days-cultured adipo-, osteo-, and chondrocytes. A volume of 175μl RPMI-1640 Dutch Modified medium containing 0.875×10^6^ primary BCP-ALL cells (1.0×10^6^ cells/ml) was added either directly to the well or in a transwell (0.4μm pore-size; Corning, New York, USA). The MSCs, adipo-, osteo-, chondrocytes, or fibroblasts and BCP-ALL cells were mono- or co-cultured for 5 days. Supernatant was collected as described previously to detect cyto-/chemokine secretion levels by Luminex technology. Cells were harvested and stained with antibodies distinguishing BCP-ALL cells from MSCs, fibroblasts, adipo-, osteo-, or chondrocytes, which included Brilliant violet 421 anti-human CD19 (Biolegend), and Alexa Fluor 750-conjugated human ALCAM/CD166 (R&D Systems, Minneapolis, MN, USA). Sytox Red Dead Cell Stain (ThermoFisher Scientific) was used to select for the viable population. Cells were measured by flow cytometry using a Cytoflex S (Beckman&Coulter, Brea, CA, USA). Flow cytometry data were analyzed by using FlowJo (version 10.7.1) and graphs were created by GraphPad Prism (version 9.1.2).

### Quantification of secreted cytokine and chemokine levels

An in-house developed and validated multiplex immunoassay based on Luminex technology (xMAP, Luminex, Austin, TX, USA) was used to quantify secreted levels of cyto-/chemokines by the Laboratory of Translational Immunology (UMC Utrecht, the Netherlands) ^22^. Samples were incubated with antibody-conjugated MagPlex microspheres for 1 hour at room temperature with constant shaking. Samples were incubated 1 hour with biotinylated antibodies, followed by 10 minutes of incubation with phycoerythrin (PE)-conjugated streptavidin. PE-conjugated streptavidin was diluted in high performance ELISA buffer (HPE) (Sanquin, Amsterdam, The Netherlands). The Biorad FlexMAP3D (Biorad laboratories, Hercules, CA, USA) and xPONENT software version 4.2 (Luminex) was used to determine the concentration of cyto-/chemokines in collected supernatants and plain culture media using standard curves per analyte. Bio-Plex Manager software, version 6.1.1 (BioRad) was used to analyze the data. T-SNE plot was created using R (version 4.1.3, Rtsne package), perplexity value 8, to identify clusters characterized by (dis)similarities in cyto-/chemokine secretion profiles. GraphPad Prism (version 9.1.2) was used to create graphs.

### Dye transfer experiments

Vybrant DiI Cell-Labeling Solution (Invitrogen) was used to pre-stain primary BCP-ALL cells for 4 hours. DiI-stained ALL cells (0.875×10^6^) were added to MSCs, adipo-, osteo-, chondrocytes, or fibroblasts in 24-well plates and cultured for 16 hours at 37°C/5%CO_2_. Cells were harvested and stained with antibodies as indicated previously. Dye transfer from leukemic cells to supportive tissues was visualized by excitation with 561nm laser and emission detection with 585/42 filter (Cytoflex S).

### Statistical analysis

One-way ANOVA tests and one or two-sided (un)paired t-tests were performed using GraphPad Prism (version 9.1.2), and correction for multiple testing was included when appropriate. A p-value below 0.05 was considered statistically significant.

## Results

### The origin of MSCs does not affect the survival benefit gained by leukemic cells

Leukemic cells from 19 different BCP-ALL patients were co-cultured with MSCs obtained at diagnosis of BCP-ALL (n=4), with MSCs collected at the end of consolidation treatment of the same patients (timepoint day 79, n=4) and with MSCs collected from healthy, non-leukemic donors (n=2; ***Supplementary table 4***). Co-culture of leukemic cells with diagnosis-MSCs (day 0) and consolidation-MSCs (day 79) resulted in similar survival benefit for leukemic cells (p=0.069, **Figure 1A**). MSCs from two non-leukemic donors provided similar benefit as MSCs from leukemia patients (normal vs. day 0: p=0.225, normal vs. day 79: p=0.106, **Figure 1A**). These data show that the timepoint of collection (diagnosis/full blown leukemia versus in remission/end of consolidation therapy) and the origin of the MSCs (leukemia patient or healthy control) does not differentially affect the survival benefit for BCP-ALL cells. This also indicates that different origins of primary MSCs can be used for *ex vivo* BCP-ALL studies.

**Figure 1.**
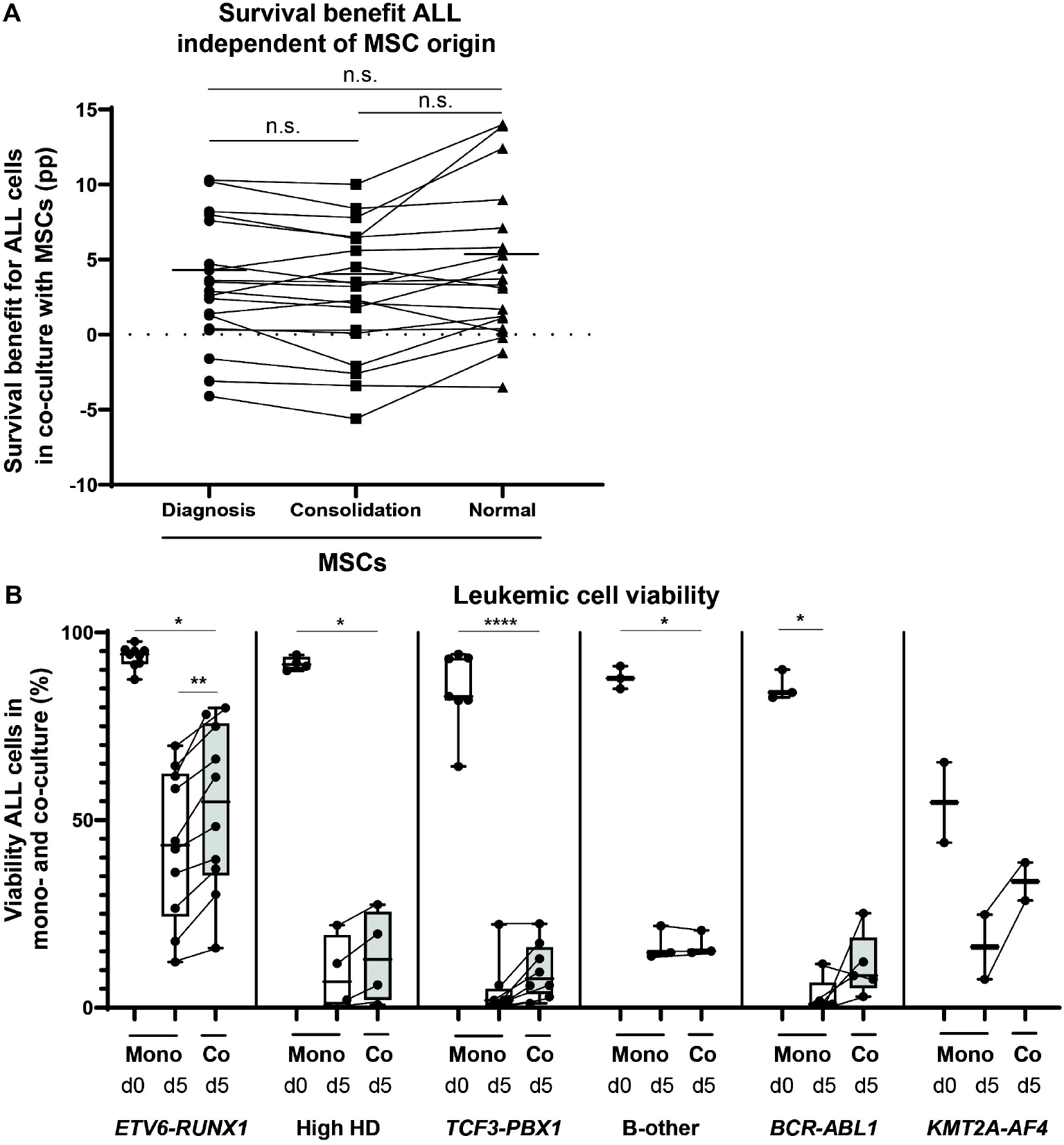
MSC-induced survival benefit for leukemic cells is not affected by the origin of MSCs but differs between subtypes of BCP-ALL. **(A)** Y-axis represents the percent point (pp) survival benefit for patients’ leukemic cells upon co-culture with paired MSCs isolated from bone marrow samples taken at diagnosis of ALL (circles) and after consolidation therapy (squares), or those taken from healthy controls (normal; triangles) compared with the survival of ALL cells kept in mono-culture. Each of 19 ALL cases were co-cultured with 4 diagnosis-MSCs (day 0), 4 consolidation-MSCs (day 79), and 2 normal-MSCs; these biological replicates were averaged. Dotted line indicates no effect on the survival of BCP-ALL cells in co-culture compared to mono-culture. Median benefit is indicated by solid line. n.s. indicates not significant. **(B)** Percentage of viable cells in ALL mono-culture (white boxplots) on day 0 (after thawing) and after 5 days of mono-culture. The percentage of viability in co-cultures of the same ALL and MSCs after 5 days of culture is depicted in the grey boxplots. The average of four independent MSC co-cultures is shown (MSC#1, #3, #6 and #7). Paired samples are connected by lines. ALL samples include *ETV6-RUNX1* (n=10), high hyperdiploid (HD) (n=4), *TCF3-PBX1* (n=8), B-other (n=3), *BCR-ABL1* (n=5), and *KMT2A-AF4* (n=2). * p < 0.05, ** p < 0.01, **** p < 0.0001.

### MSC-induced survival benefit differs between BCP-ALL subtypes

The viability of BCP-ALL patients’ cells in *ex vivo* assays rapidly diminishes if these cells are kept in mono-culture, e.g., viability of BCP-ALL cells decreases from 83%±14.3 upon recovery after liquid nitrogen storage to only 15.5%±14.8 after 5 days of mono-culture (**Figure 1B**). The most pronounced reduction in viability is observed for *TCF3-PBX1* and *BCR-ABL1*-positive ALL cases, exemplified by the fact that the percentage of viable cells for 6 out of 8 *TCF3-PBX1* and 4 out of 5 *BCR-ABL1* positive cases was severely reduced to a median of less than 3% after 5 days of mono-culture compared with initial viability of median 83% (*TCF3-PBX1*; range 64.3-94.2%) and 84% (*BCR-ABL1*; range 82.7-90.1%) at the start of *ex vivo* mono-cultures. Co-culture of BCP-ALL cells with MSCs provided a survival benefit to most cases: a higher viability was seen in co-culture compared with mono-culture for *BCR-ALB1* (8.6-fold) and *TCF3-PBX1* (3.9-fold) ALL cases, although not significant. On average the viability in co-culture was higher than in mono-culture for 75% of the BCP-ALL cases (**Figure 1B**). Although limited to 2 samples, the median survival benefit for *KMT2A-AF4* positive ALL cells clearly increased by co-culture with MSCs (16.2% in mono- to 33.7% in co-culture). *ETV6-RUNX1* positive cases showed a relatively high median viability in mono-culture (43.4%) compared with other subtypes and this viability increased even more by stromal support (54.9%; 1.3-fold, p=0.025) (**Figure 1B; *Supplementary table 4***). These results indicate that survival benefit is provided independent of the MSC origin, but that the need for and benefit from stromal support differs between genetic subtypes of BCP-ALL.

### Different supportive cell types give a survival benefit to BCP-ALL cells

MSCs collected from patients remained multipotent as exemplified by their potency to differentiate into adipo-, osteo-, and chondrocytes (**Figure 2A-C**). A gradual loss of differentiation potential for MSCs was evident from passage 12 onwards (***Supplementary table 3***). We therefore use MSCs cultured for a maximum of 10 passages for our subsequent studies.

**Figure 2.**
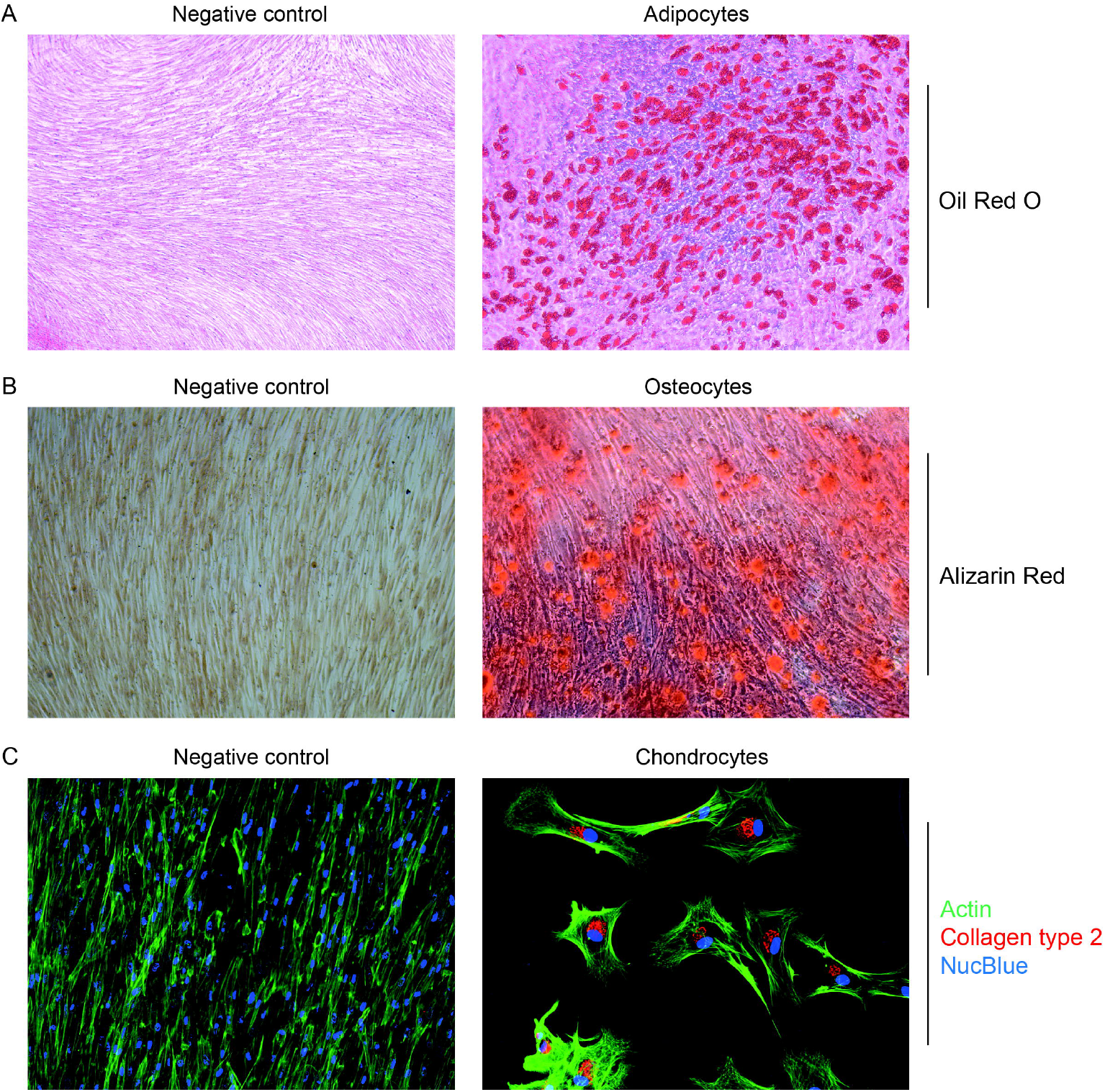
Bone marrow-derived MSCs have multi-lineage potential. A representative experiment is depicted in which multi-lineage potential of MSC#8 into **(A)** adipocytes, **(B)** osteocytes, and **(C)** chondrocytes was confirmed after 21 days of culture. Differentiation into the mesodermal lineages was verified by Oil Red O, Alizarin Red, Actin/NucBlue/Collagen type-2 antibody staining, respectively. Images of adipo- and osteocytes were captured with a 5X and 10X magnification, resp.; chondrocytes with a 20X magnification.

Four BCP-ALL patient samples were co-cultured with distinct supportive cell types found in the bone marrow: MSCs, adipocytes, osteocytes, chondrocytes, and fibroblasts. The median viability of cells from these 4 cases in mono-culture was 49.5% and increased when kept in co-culture with supportive cell types (**Figure 3A**). A significant increase in survival of patients’ derived leukemic cells in co-cultures was observed for MSCs (median +33.3pp, p=0.0112), osteocytes (+29.9pp, p=0.0081), and fibroblasts (+35.6pp, p=0.0002) (**Figure 3B**). The viability of BCP-ALL samples also increased, albeit not statistically significant in co-culture with chondrocytes (+17.9pp, p=0.1383). No or only limited benefit was provided by fully differentiated adipocytes (<5pp; **Figure 3A/B**).

**Figure 3.**
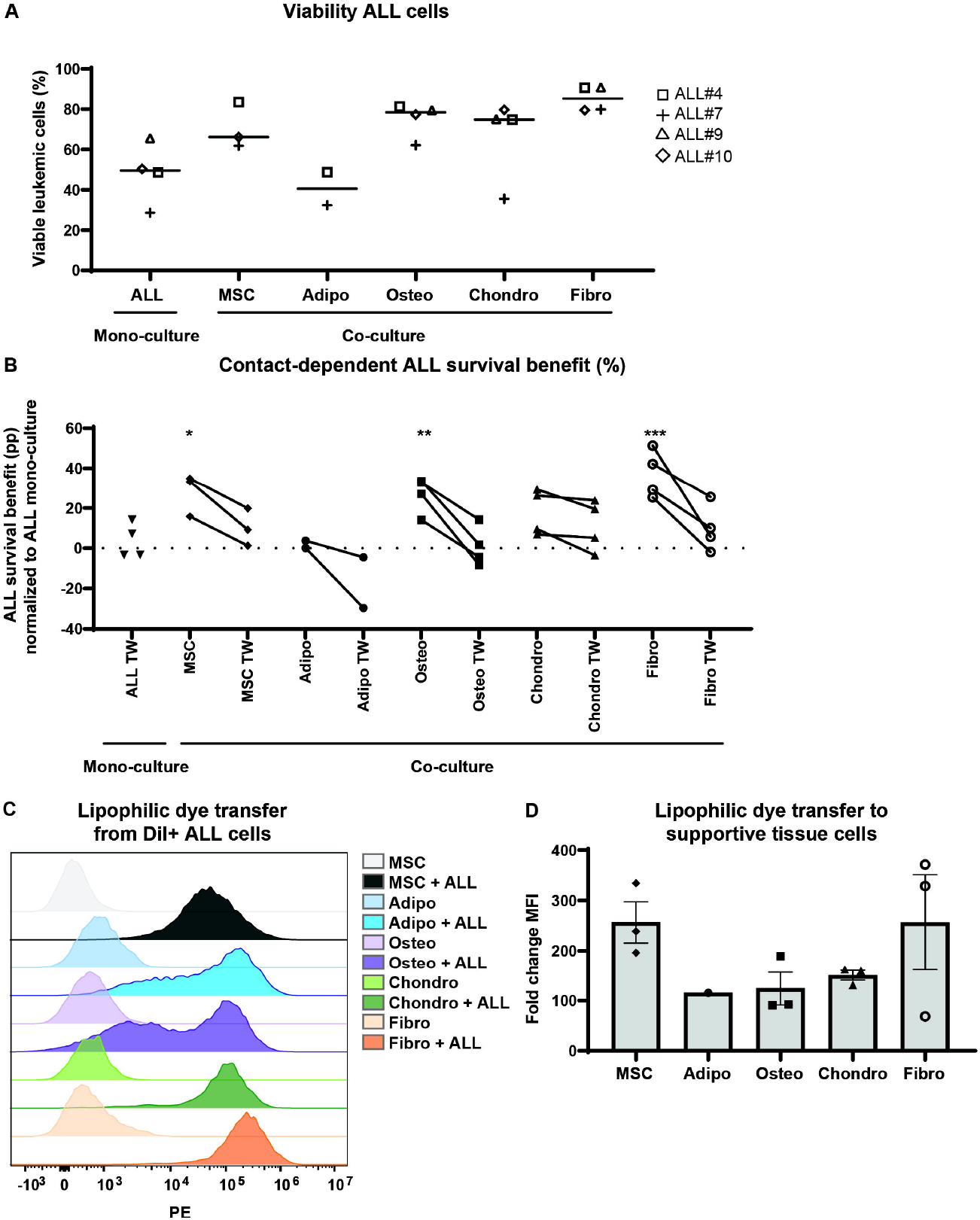
Patient’s leukemic cell survival benefit induced by supportive tissue cells is contact-dependent. **(A)** Y-axis represents the viability of ALL cells upon mono-culture or co-culture with MSCs (MSC#8), adipocytes, osteocytes, chondrocytes, or fibroblasts. Open square, plus, open triangle, and open diamond indicate ALL#4, ALL#7, ALL#9, and ALL#10, resp. **(B)** Y-axis depicts the percent point (pp) survival benefit for ALL cells cultured directly or indirectly (0.4mm-pore size transwell (TW) setting) with MSCs (diamonds), adipocytes (closed circles), osteocytes (squares), chondrocytes (triangles), and fibroblasts (open circles). Paired samples are connected by lines. At the left, indicated by triangles, the survival benefit of ALL cells cultured in the insert of a transwell setting has been compared to the survival of ALL cells kept in normal mono-culture, which is similar to culture in the bottom compartment of a transwell system. ALL#4, ALL#7, ALL#9, and ALL#10 were used for these co-cultures. Technical triplicates were averaged for 4 (osteo-/chondrocytes and fibroblasts), 3 (MSCs), and 2 (adipocytes) independent experiments. Dotted line indicates no effect on the survival of BCP-ALL cells in co-culture compared to mono-culture. **(C)** A representative experiment is depicted for DiI dye transfer to MSCs (black), adipocytes (blue), osteocytes (purple), chondrocytes (green), or fibroblasts (orange) upon co-culture with DiI-positive primary leukemic cells (ALL#7, #9 and #10) for 16 hours as determined by flow cytometry. **(D)** Fold change of the mean fluorescent intensity (MFI) in the PE-channel was quantified for each cell type by dividing the MFI of supportive tissue cells kept in co-culture with the DiI-labeled leukemic cells by the MFI of the corresponding tissue in mono-culture. Bars represent the average ± SEM of technical triplicates for 3 independent experiments; only dye transfer data from mature adipocytes were included (n=1). * p < 0.05, *** p < 0.001.

### Stromal support depends on direct cell-cell contact

Lipophilic DiI dye-transfer from pre-loaded BCP-ALL cells occurred to all supportive tissue types (**Figure 3C/D** and ***Supplementary Figure 1***), which is indicative of functional tunneling nanotubes formation that are involved in direct intercellular exchange of various molecules and increased viability of leukemic cells ^15^. Disruption of direct cell-cell contact in a transwell setting reduced the survival benefit provided by the various supportive tissues towards the viability seen when leukemic cells were kept in mono-culture (**Figure 3B**). This decrease in viability of disconnected BCP-ALL cells could not be attributed to a parallel reduction in the number of supportive tissue cells (***Supplementary Figure 2***).

### Cyto-/chemokine profile of supportive tissue cells and BCP-ALL cells

T-SNE analysis of 171 cyto-/chemokines that were secreted in the supernatant of direct co-cultures of BCP-ALL and MSCs compared with those released by (the sum of) BCP-ALL and MSCs kept in mono-culture revealed a change in the secretome (***Supplementary Figure 3***). Physical separation of MSCs and BCP-ALL cells using a transwell setting decreased secretion of several cyto-/chemokines compared with direct co-culture (***Supplementary Figure 4***). Eight cyto-/chemokines for which the secreted levels increased in the co-culture setting of BCP-ALL cells and MSCs, were also analyzed in the other types of supportive tissues (**Figure 4A**). The concentration of secreted cyto-/chemokines differed between mono-cultures of supportive tissue types, e.g., osteocytes secreted higher levels of CXCL1, CXCL5, CXCL8, and CCL2 compared with MSCs, whereas MSCs secreted higher levels of IL6, and CXCL10 compared with the other supportive tissue types. In contrast to MSCs and the other supportive tissues, BCP-ALL cells hardly produced cyto-/chemokines (**Figure 4B**). Direct co-cultures of BCP-ALL cells and supportive cells most often resulted in a further increase of secreted cyto-/chemokine levels compared with the sum of the corresponding mono-cultures. CCL22 secretion was increased in BCP-ALL/MSC co-cultures (1.41-fold, p=0.001). Secreted levels were also higher after co-culture with osteocytes and fibroblasts, while decreased CCL22 secretion was observed in co-culture with adipocytes and chondrocytes. The most consistent raise in secreted levels imposed by ALL cells was observed for CXCL10 (1.39-fold, p=0.003), IL6 (1.29-fold, p=0.013), and CXCL5 (1.18-fold, p=0.006), which was observed for all tested supportive tissue types whereas the secreted levels by ALL cells in mono-culture was low (**Figure 4B**; ***Supplementary Figure 5***). The increase in cyto-/chemokine levels was not caused by an increased number of supportive cells, since these cell numbers remained similar in mono- and co-culture (***Supplementary Figure 2***). This indicates that secretion of cyto-/chemokines is induced by BCP-ALL cells upon co-culture with the supportive tissues.

**Figure 4.**
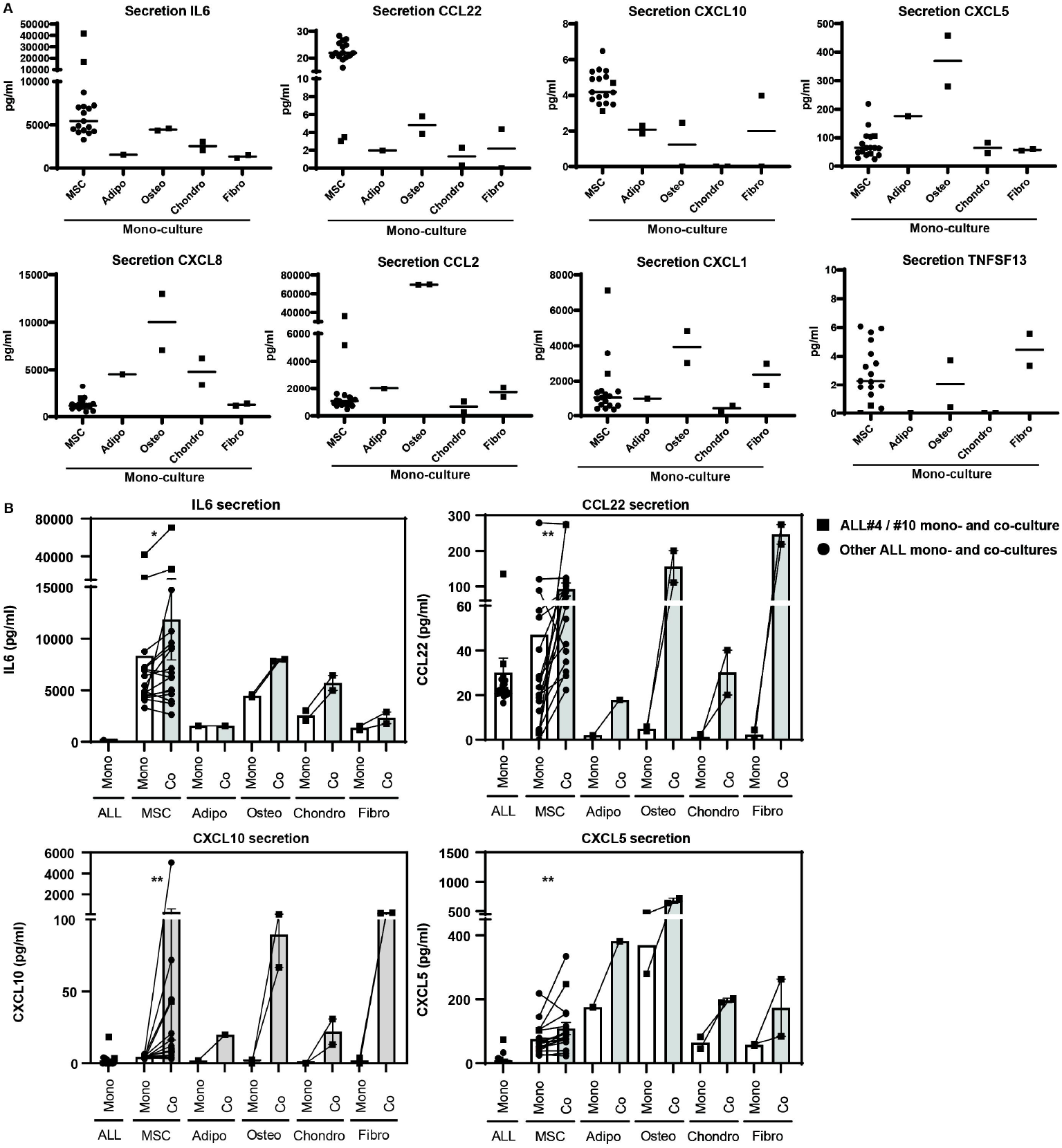
BCP-ALL cells induce cytokine secretion upon co-culture with multiple supportive tissue types. **(A)** Graphs depict median secretion level (pg/ml) of IL6, CCL22, CXCL10, CXCL5, CXCL8, CCL2, CXCL1, and TNFSF13 in mono-cultures of MSCs (n=17), adipocytes (n=1), osteocytes (n=2), chondrocytes (n=2), or fibroblasts (n=2). **(B)** Graphs depict the secreted level (pg/ml) of IL6, CCL22, CXCL10, and CXCL5 in mono-(white) and co-cultures (grey) of patients’ BCP-ALL cells and the supportive tissue types. Supportive tissue mono-cultures are connected by lines to the levels measured in co-cultures of the same samples. Squares indicate mono- and co-culture of ALL#4 and ALL#10. Other mono- and co-cultures are indicated with circles (ALL#6-7, ALL#33-44). * p < 0.05, ** p < 0.01.

## Discussion

We show that the viability of patients’ BCP-ALL cells benefits from co-culture with different origins of MSCs (i.e., those collected at time of leukemia, at remission or from healthy controls) and types of supportive bone marrow cells. The need for this support clearly differed between subtypes of BCP-ALL, with *BCR-ABL1*-positive and *TCF3-PBX1*-positive subtypes being the most dependent on stromal tissue support. In addition, the secretion of several cyto-/chemokines (which most often are produced by the supportive tissues) increased when these cells were exposed to BCP-ALL cells. This underlines our earlier finding by De Rooij *et al*. ^23^, in which the cyto-/chemokine secretion profile was determined by the subtype of ALL and not by the origin of MSCs that was used. These data suggests that leukemic cells drive the changes in the bone marrow microenvironment, a phenomenon we refer to as the ALL-educated niche.

We noticed that mature adipocytes did not affect the viability of BCP-ALL cells nor induced major changes in the secretion of cyto-/chemokines, suggesting that adipocytes, albeit a major component of the bone marrow, are less detrimental in the support of leukemic cells. For the other supportive tissues this may be different since beside MSCs also osteocytes, fibroblasts, and to a lesser extend chondrocytes are supportive to leukemic cells in a contact-dependent way. Chondrocytes are known to secrete growth factors regulating synthesis of the extracellular matrix (ECM) ^24^. In case of cancer, the ECM becomes abnormal which can deregulate stromal cell behavior and stimulate inflammation, leading to a tumor-facilitating microenvironment ^25^. In acute myeloid leukemia (AML), crosstalk between osteoblasts and the leukemic cells results in local cytokine release that stimulates the proliferation of AML cells ^26^. In ALL, we observed that an interferon (IFN)-related gene expression profile was induced by BCP-ALL cells in MSCs, including CXCL10 expression ^27^. This signature was partly dependent on cross-talk via tunneling nanotubes (TNTs) as the expression of several IFN-related genes decreased when both cell types were physically disconnected which coincided with reduced lipophilic dye transfer as previously demonstrated by our group ^15^. In line with an ALL-educated niche caused by direct cell-cell communication, Burt *et al*. showed that transfer of mitochondria derived from cancer-associated fibroblasts (CAFs) protected ALL cells against chemotherapy-induced reactive oxygen species (ROS) and apoptosis ^28^. The CAF-mediated protection was abrogated when the physical connection between both cell types was blocked ^28^. The transport of mitochondria may be bi-directional since we also observed that mitochondria can be transported from BCP-ALL cells to MSCs ^29^. Mitochondrial reactive oxygen species (mtROS) influence the production of cytokines, amongst others IL6 ^30,31^.

IL6 is important for monocyte differentiation into macrophages, B-cell differentiation into antibody producing plasma cells, and the proliferation of hematopoietic stem cells ^32^. IL6 can also exert pro-tumorigenic activities such as supporting proliferation of cancer cells, and evasion of immune surveillance ^32,33^. Interestingly, high IL6 levels were shown in bone marrow samples of BCP-ALL patients at the time of diagnosis, while these cytokine levels were significantly decreased after induction therapy ^33^. Despite a limited sample size, we noticed a simultaneous decrease in IL6 and CCL22 secretion and in the survival benefit provided by MSCs when the direct, cell-cell interaction between leukemic cells and MSCs was prevented.

In contrast to IL6, the CXCL-family and CCL2/CCL22 are more involved in recruitment and accumulation of (suppressive) immune cells. CCL22 recruits regulatory T-cells into tumor tissues ^34^. CCL2-attracted monocytes transform into immunosuppressive macrophages ^35^. CXCL5 recruits neutrophils and promotes infiltration of myeloid-derived suppressor cells in the tumor microenvironment ^36^. These cyto-/chemokines are produced by a.o. MSCs (this study) and have been shown to induce an immune-suppressive environment in the vicinity of cancer cells ^37,38^. Together these findings strengthen our earlier observation that BCP-ALL cells hijack the bone marrow microenvironment and offers the perspective that interference with this stromal interaction and/or released cyto-/chemokines may be of additive value in the treatment of BCP-ALL.

## Supporting information

Supplemental Tabels

## Acknowledgements

We would like to thank all group members of the Erasmus Medical Center in Rotterdam, and Princess Máxima Center for Pediatric Oncology in Utrecht for their project input and help in (leukemic) sample processing. We thank Prof. dr. Gerjo van Osch from the Erasmus Medical Center in Rotterdam for her advice on the HDFα cell line. We also thank the MultiPlex Core Facility in the Center for Translational Immunology in Utrecht for performing Luminex measurements (Stefan Nierkens). This study was funded by the Pediatric Oncology Foundation Rotterdam (SKOCR) and the Oncode Institute in Utrecht.

## Contributors

MS and CvdV contributed to study design, performed the experiments, collected data, performed data analysis, and wrote the paper for this study. MLdB designed the study and finalized this manuscript. JO, AB, and CEJR performed experiments. JB performed statistical analysis. All authors reviewed and approved the final manuscript.

## References

1. Pui CH, Carroll WL, Meshinchi S, Arceci RJ. Biology, risk stratification, and therapy of pediatric acute leukemias: An update. J Clin Oncol. 2011;29(5):551–565. doi:10.1200/JCO.2010.30.7405

2. Kato M, Manabe A. Treatment and biology of pediatric acute lymphoblastic leukemia. Pediatr Int. 2018;60(1):4–12. doi:10.1111/ped.13457

3. Li JF, Ma XJ, Ying LL, Tong YH, Xiang XP. Multi-Omics Analysis of Acute Lymphoblastic Leukemia Identified the Methylation and Expression Differences Between BCP-ALL and T-ALL. Front Cell Dev Biol. 2021;8(January):1–8. doi:10.3389/fcell.2020.622393

4. Boer JM, Steeghs EMP, Marchante JRM, et al. Tyrosine kinase fusion genes in pediatric BCR-ABL1-like acute lymphoblastic leukemia. Oncotarget. 2017;8(3):4618–4628. doi:10.18632/oncotarget.13492

5. Hunger SP, Mullighan CG. Acute lymphoblastic leukemia in children. N Engl J Med. 2015;373(16):1541–1552. doi:10.1056/NEJMra1400972

6. Teachey DT, Pui CH. Comparative features and outcomes between paediatric T-cell and B-cell acute lymphoblastic leukaemia. Lancet Oncol. 2019;20(3):e142–e154. doi:10.1016/S1470-2045(19)30031-2

7. Den Boer ML, van Slegtenhorst M, De Menezes RX, et al. A subtype of childhood acute lymphoblastic leukaemia with poor treatment outcome: a genome-wide classification study. Lancet Oncol. 2009;10(2):125–134. doi:10.1016/S1470-2045(08)70339-5

8. Pieters R, De Groot-Kruseman H, Van Der Velden V, et al. Successful therapy reduction and intensification for childhood acute lymphoblastic leukemia based on minimal residual disease monitoring: Study ALL10 from the Dutch Childhood Oncology Group. J Clin Oncol. 2016;34(22):2591–2601. doi:10.1200/JCO.2015.64.6364

9. Schwab C, Harrison CJ. Advances in B-cell Precursor Acute Lymphoblastic Leukemia Genomics. HemaSphere. Published online 2018:1. doi:10.1097/hs9.0000000000000053

10. Naderi EH, Skah S, Ugland H, et al. Bone marrow stroma-derived PGE2 protects BCP-ALL cells from DNA damage-induced p53 accumulation and cell death. Mol Cancer. 2015;14(1):1–12. doi:10.1186/s12943-014-0278-9

11. Hong IS, Lee HY, Kang KS. Mesenchymal stem cells and cancer: Friends or enemies? Mutat Res - Fundam Mol Mech Mutagen. 2014;768(C):98–106. doi:10.1016/j.mrfmmm.2014.01.006

12. Usmani S, Sivagnanalingam U, Tkachenko O, Nunez L, Shand JC, Mullen CA. Support of acute lymphoblastic leukemia cells by nonmalignant bone marrow stromal cells. Oncol Lett. 2019;17(6):5039–5049. doi:10.3892/ol.2019.10188

13. Duan CW, Shi J, Chen J, et al. Leukemia propagating cells rebuild an evolving niche in response to therapy. Cancer Cell. 2014;25(6):778–793. doi:10.1016/j.ccr.2014.04.015

14. Portale F, Cricrì G, Bresolin S, et al. ActivinA: a new leukemia-promoting factor conferring migratory advantage to B-cell precursor-acute lymphoblastic leukemic cells. Haematologica. 2019;104(3):533–545. doi:10.3324/haematol.2018.188664

15. Polak R, De Rooij B, Pieters R, Den Boer ML. B-cell precursor acute lymphoblastic leukemia cells use tunneling nanotubes to orchestrate their microenvironment. Blood. 2015;126(21):2404–2414. doi:10.1182/blood-2015-03-634238

16. Tamma R, Ribatti D. Bone niches, hematopoietic stem cells, and vessel formation. Int J Mol Sci. 2017;18(1). doi:10.3390/ijms18010151

17. Dander E, Palmi C, D’amico G, Cazzaniga G. The bone marrow niche in b-cell acute lymphoblastic leukemia: The role of microenvironment from pre-leukemia to overt leukemia. Int J Mol Sci. 2021;22(9). doi:10.3390/ijms22094426

18. Pastorczak A, Domka K, Fidyt K, Poprzeczko M, Firczuk M. Lymphoblastic Leukemia. Published online 2021:1–25.

19. Baryawno N, Przybylski D, Kowalczyk MS, et al. A Cellular Taxonomy of the Bone Marrow Stroma in Homeostasis and Leukemia. Cell. 2019;177(7):1915–1932.e16. doi:10.1016/j.cell.2019.04.040

20. Ariës IM, Jerchel IS, Van Den Dungen RESR, et al. EMP1, a novel poor prognostic factor in pediatric leukemia regulates prednisolone resistance, cell proliferation, migration and adhesion. Leukemia. 2014;28(9):1828–1837. doi:10.1038/leu.2014.80

21. Den Boer ML, Harms DO, Pieters R, et al. Patient stratification based on prednisolone-vincristine-asparaginase resistance profiles in children with acute lymphoblastic leukemia. J Clin Oncol. 2003;21(17):3262–3268. doi:10.1200/JCO.2003.11.031

22. Smids C, Horje CSHT, Nierkens S, et al. Candidate serum markers in early Crohn’s disease: Predictors of disease course. J Crohn’s Colitis. 2017;11(9):1090–1100. doi:10.1093/ecco-jcc/jjx049

23. De Rooij B, Polak R. Acute lymphoblastic leukemia cells create a leukemic niche without affecting the CXCR4/CXCL12 axis. Haematologica. Published online 2017:389–393. http://www.haematologica.org.

24. Chen H, Tan XN, Hu S, et al. Molecular Mechanisms of Chondrocyte Proliferation and Differentiation. Front Cell Dev Biol. 2021;9(May):1–13. doi:10.3389/fcell.2021.664168

25. Lu P, Weaver VM, Werb Z. The extracellular matrix: A dynamic niche in cancer progression. J Cell Biol. 2012;196(4):395–406. doi:10.1083/jcb.201102147

26. Bruserud Ø, Reikvam H, Brenner AK. Toll-like Receptor 4, Osteoblasts and Leukemogenesis; the Lesson from Acute Myeloid Leukemia. Molecules. 2022;27(3). doi:10.3390/molecules27030735

27. Smeets MWE, Steeghs EMP, Orsel J, et al. B-cell precursor acute lymphoblastic leukemia elicits an interferon-alpha/beta response in bone marrow-derived mesenchymal stroma. bioRxiv. doi:10.1101/2023.05.09.539232

28. Burt R, Dey A, Aref S, et al. Activated stromal cells transfer mitochondria to rescue acute lymphoblastic leukemia cells from oxidative stress. Blood. 2019;134(17):1415–1429. doi:10.1182/blood.2019001398

29. De Rooij B, Polak R, Stalpers F, Pieters R, Den Boer ML. Tunneling nanotubes facilitate autophagosome transfer in the leukemic niche. Leukemia. 2017;31(7):1651–1654. doi:10.1038/leu.2017.117

30. Naik E, Dixit VM. Mitochondrial reactive oxygen species drive proinflammatory cytokine production. J Exp Med. 2011;208(3):417–420. doi:10.1084/jem.20110367

31. Herb M, Gluschko A, Wiegmann K, et al. Mitochondrial reactive oxygen species enable proinflammatory signaling through disulfide linkage of NEMO. Sci Signal. 2019;12(568). doi:10.1126/scisignal.aar5926

32. Diehl S, Rincón M. The two faces of IL-6 on Th1 / Th2 differentiation. 2002;39:531–536.

33. Magalhães-Gama F, Kerr MWA, D. Araújo ND, et al. Imbalance of Chemokines and Cytokines in the Bone Marrow Microenvironment of Children with B-Cell Acute Lymphoblastic Leukemia. J Oncol. 2021;2021. doi:10.1155/2021/5530650

34. Jafarzadeh A, Fooladseresht H, Minaee K, et al. Higher circulating levels of chemokine CCL22 in patients with breast cancer: evaluation of the influences of tumor stage and chemokine gene polymorphism. Tumor Biol. 2015;36(2):1163–1171. doi:10.1007/s13277-014-2739-6

35. Gok Yavuz B, Gunaydin G, Gedik ME, et al. Cancer associated fibroblasts sculpt tumour microenvironment by recruiting monocytes and inducing immunosuppressive PD-1 + TAMs. Sci Rep. 2019;9(1):1–15. doi:10.1038/s41598-019-39553-z

36. Zhang W, Wang H, Sun M, et al. CXCL5/CXCR2 axis in tumor microenvironment as potential diagnostic biomarker and therapeutic target. Cancer Commun. 2020;40(2-3):69–80. doi:10.1002/cac2.12010

37. Rapp M, Wintergerst MWM, Kunz WG, et al. CCL22 controls immunity by promoting regulatory T cell communication with dendritic cells in lymph nodes. J Exp Med. 2019;216(5):1170–1181. doi:10.1084/jem.20170277

38. Röhrle N., Knott M.M.L., Anz D. (2020) CCL22 Signaling in the Tumor Environment. In: Birbrair A. (eds) Tumor Microenvironment. Advances in Experimental Medicine and Biology, vol 1231. Springer, Cham. https://doi.org/10.1007/978-3-030-36667-4_8.

