## Supplemental Figures for "Various Cell Types in the Bone Marrow sustain Primary B-Cell Precursor Acute Lymphoblastic Leukemia"

### Supplementary figures

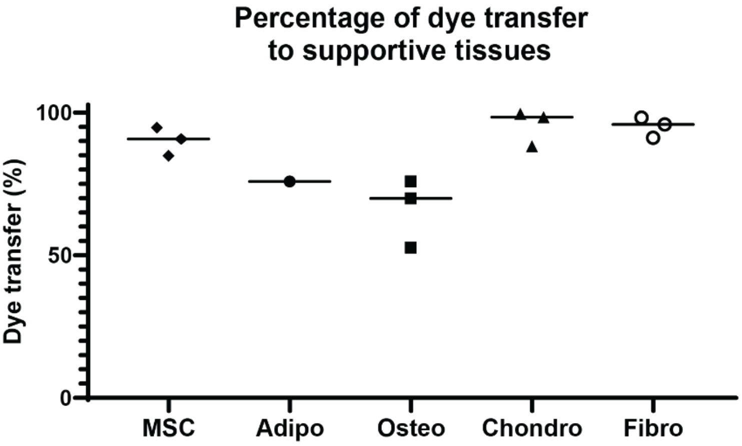


**Supplementary Figure 1. Dye transfer from primary leukemic cells to supportive tissue cells.** Graph showing the percentage of supportive cells becoming positive for the lipophilic dye derived from pre-loaded BCP-ALL cells. DiI-positive leukemic cells from 3 different patients were co-cultured with the indicated supportive tissue cells for 16 hours and the percentage of dye-positive cells was measured flow cytometry. Values represent median of 3 independent experiments, technical triplicates per experiment were averaged. Only dye transfer data from mature adipocytes were included (n=1).

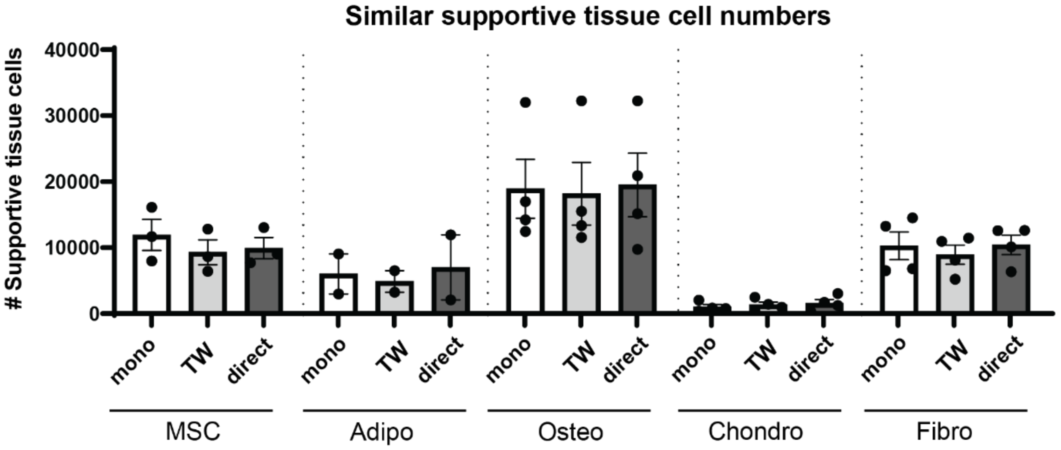


**Supplementary Figure 2. Cell numbers of supportive tissues are comparable between mono- and (transwell) co-culture**. Y-axis represents the number of supportive tissue cells in mono-culture (white), in the bottom compartment of a 0.4µm-pore size transwell (TW) setting upon BCP-ALL co-culture (grey), or upon direct BCP-ALL co-culture (dark grey). Values represent mean ± SEM of 4 (osteocytes, chondrocytes, and fibroblasts), 3 (MSCs) or 2 (adipocytes) independent experiments, technical triplicates per experiment were averaged. Only data from mature adipocytes were included.

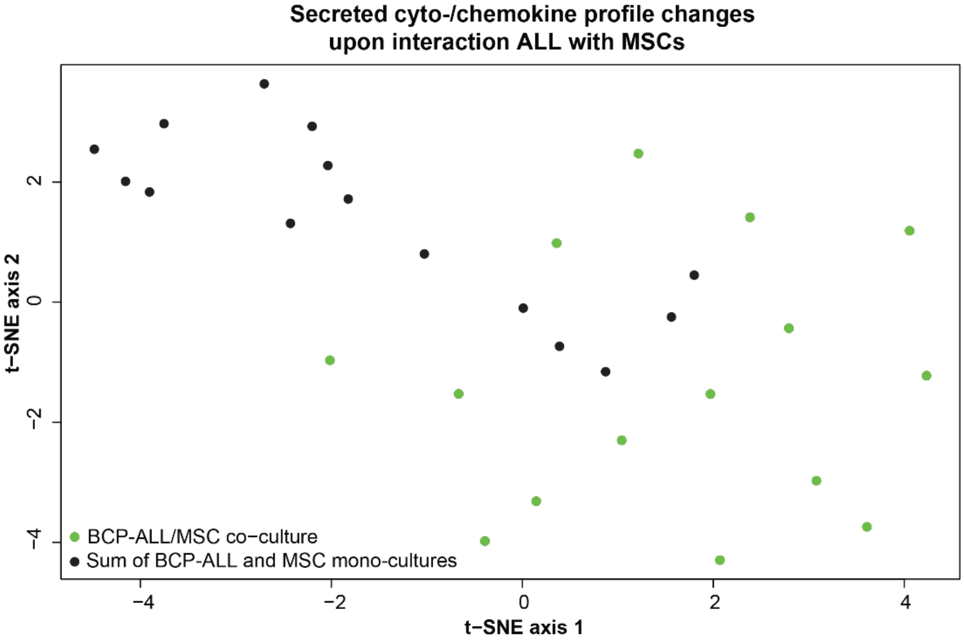


**Supplementary Figure 3. The secreted cyto-/chemokine profile alters upon BCP-ALL/MSC co-culture.** T-SNE of secreted levels of 171 cyto-/chemokines in the sum of the monocultures (black circles) and co-cultures of BCP-ALL and MSCs (green circles).

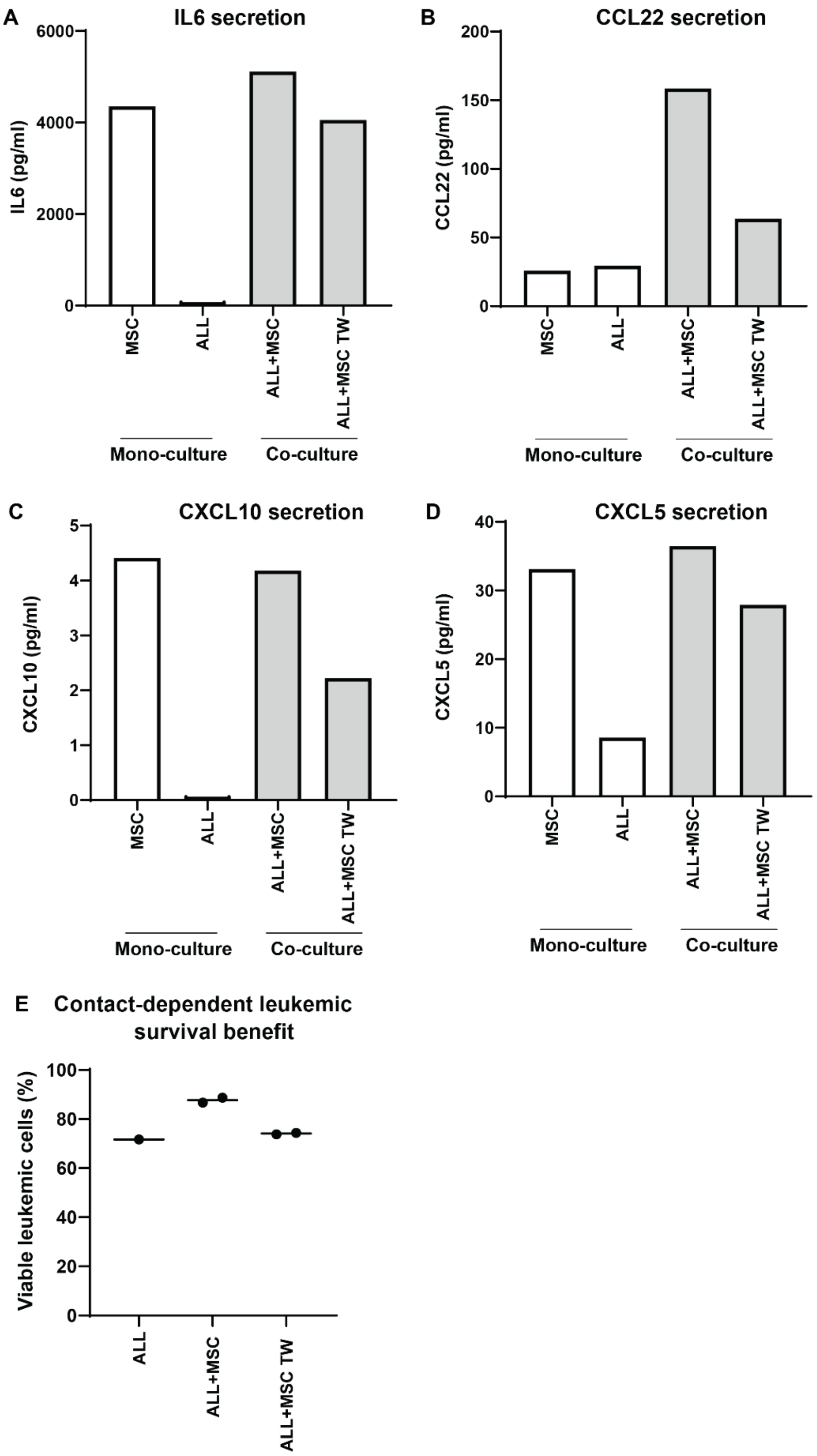


**Supplementary Figure 4. Cyto-/chemokine secretion induced by BCP-ALL cells is dependent on direct cell-cell contact**. Graphs depict the secreted level (pg/ml) of **(A)** IL6, **(B)** CCL22, **(C)** CXCL10, and **(D)** CXCL5 in mono-cultures of MSCs and patients' BCP-ALL cells (ALL#8) (white), and in direct or indirect (0.4µm-pore size transwell setting; TW) co-cultures of MSCs and BCP-ALL cells (grey). Bars represent values of one experiment. **(E)** Y-axis represents the viability of ALL cells (ALL#8) upon mono-culture and direct or indirect (TW) co-culture with MSCs (MSC#8). Values represent median of technical duplicates for one experiment.

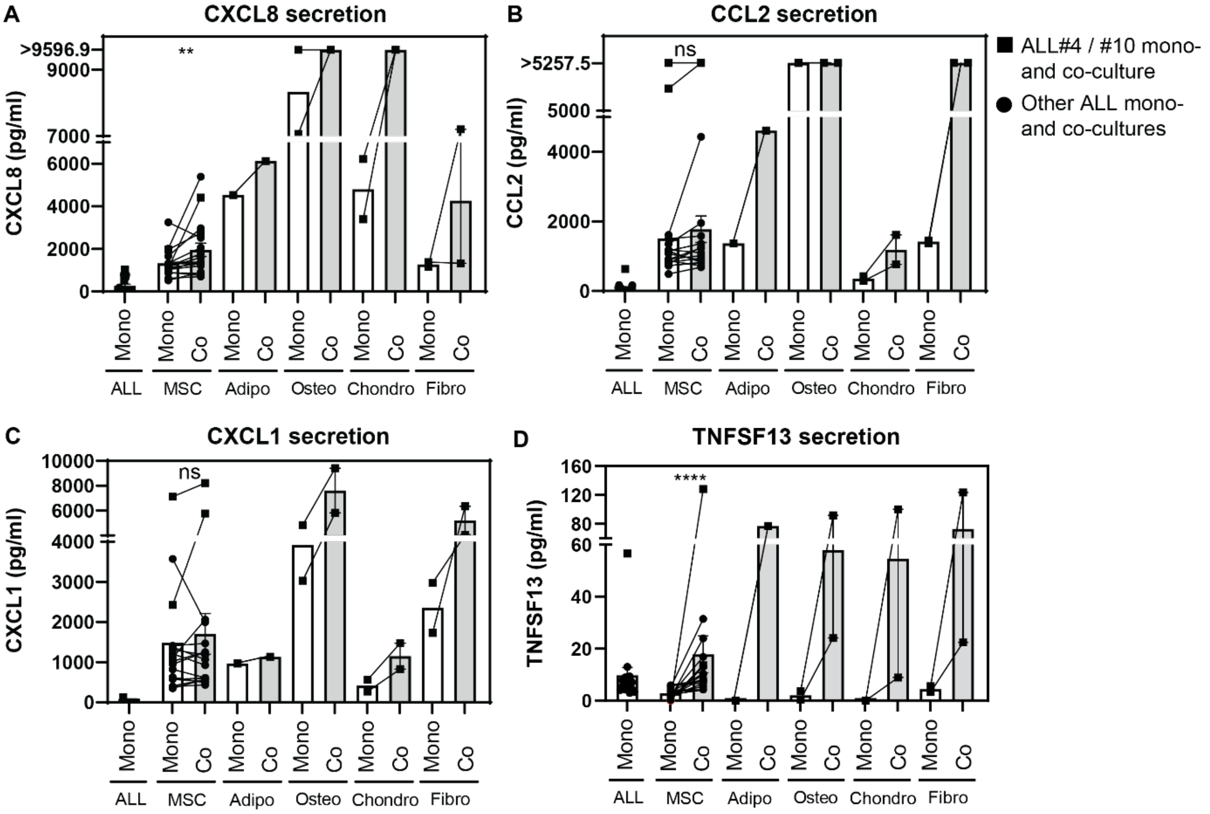


**Supplementary Figure 5. BCP-ALL cells induce cytokine secretion upon co-culture with multiple supportive tissue types.** Graphs depict the secreted level (pg/ml) of **(A)** CXCL8, **(B)** CCL2, **(C)** CXCL1, and **(D)** TNFSF13 in mono- (white) and co-cultures (grey) of patients' BCP-ALL cells and the supportive tissue types. Supportive tissue mono-cultures are connected by lines to the levels measured in co-cultures of the same samples. Squares indicate mono- and co-culture of ALL#4 and ALL#10. Other mono- and co-cultures are indicated with circles (ALL#6-7, ALL#33-44). ns indicates not significant, ** $p < 0.01$, **** $p < 0.0001$.
