## Supplemental Tabels for "Various Cell Types in the Bone Marrow sustain Primary B-Cell Precursor Acute Lymphoblastic Leukemia"

**Supplementary tables**

**Supplementary table 1. Characteristics of bone marrow-derived mesenchymal stromal cells**

| MSC # | Derived from | ALL subtype | Remark |
| --- | --- | --- | --- |
| MSC#1 | Leukemic bone marrow | B-other | Diagnosis (day 0) |
|  | Leukemic bone marrow | B-other | After consolidation therapy (day 79) |
| MSC#2 | Leukemic bone marrow | B-other | Diagnosis (day 0) |
|  | Leukemic bone marrow | B-other | After consolidation therapy (day 79) |
| MSC#3 | Leukemic bone marrow | B-other | Diagnosis (day 0) |
|  | Leukemic bone marrow | B-other | After consolidation therapy (day 79) |
| MSC#4 | Leukemic bone marrow | *ETV6-RUNX1* | Diagnosis (day 0) |
|  | Leukemic bone marrow | *ETV6-RUNX1* | After consolidation therapy (day 79) |
| MSC#5 | Leukemic bone marrow | *ETV6-RUNX1* | Diagnosis (day 0) |
|  | Leukemic bone marrow | *ETV6-RUNX1* | After consolidation therapy (day 79) |
| MSC#6 | Healthy bone marrow | - | Normal |
| MSC#7 | Healthy bone marrow | - | Normal |
| MSC#8 | Leukemic bone marrow | Hyperdiploid | Relapse |
| MSC#9 | Leukemic bone marrow | *MLL-AF4* | Relapse |
| MSC#10 | Leukemic bone marrow | T-ALL | Diagnosis (day 0) |
| MSC#11 | Leukemic bone marrow | *ETV6-RUNX1* | Relapse |
| MSC#12 | Leukemic bone marrow | *ETV6-RUNX1* | Diagnosis (day 0) |

**Supplementary table 2. Characteristics of BCP-ALL samples.** a,b Two independent cultures with the same patient cells. Cultures with the same patient cells are treated as unique samples.

| ALL# | ALL subtype | Used in figure: | ALL# | ALL subtype | Used in figure: |
| --- | --- | --- | --- | --- | --- |
| ALL#1 | *ETV6-RUNX1*^a^ | 1B | **ALL#23** | B-other (*BCR-ABL1*-like; PAX5-amplification) | 1A/B |
| ALL#2 | *ETV6-RUNX1* ^a^ | 1B | **ALL#24** | B-other | 1A/B |
| ALL#3 | *ETV6-RUNX1* | 1B | **ALL#25** | B-other | 1A/B |
| ALL#4 | *ETV6-RUNX1* | 1A/B, 3A/B, 4A/B | **ALL#26** | *BCR-ABL1* | 1B |
| ALL#5 | *ETV6-RUNX1* | 1B | **ALL#27** | *BCR-ABL1* | 1B |
| ALL#6 | *ETV6-RUNX1* | 1A/B, 4A/B | **ALL#28** | *BCR-ABL1* | 1B |
| ALL#7 | *ETV6-RUNX1* | 1B, 3A-D, 4A/B | **ALL#29** | *BCR-ABL1* | 1A/B |
| ALL#8 | *ETV6-RUNX1* | 1A/B | **ALL#30** | *BCR-ABL1* | 1A/B |
| ALL#9 | *ETV6-RUNX1* | 1A/B, 3A/B/D | **ALL#31** | *KMT2A-AF4* | 1B |
| ALL#10 | *ETV6-RUNX1* | 1A/B, 3A/B/D | **ALL#32** | *KMT2A-AF4* | 1B |
| ALL#11 | Hyperdiploid | 1A/B | **ALL#33** | *ETV6-RUNX1*-like | 4A/B |
| ALL#12 | Hyperdiploid | 1A/B | **ALL#34** | B-other | 4A/B |
| ALL#13 | Hyperdiploid | 1A/B | **ALL#35** | B-other | 4A/B |
| ALL#14 | Hyperdiploid | 1A/B | **ALL#36** | *ETV6-RUNX1* | 4A/B |
| ALL#15 | *TCF3-PBX1* | 1A/B | **ALL#37** | *ETV6-RUNX1* | 4A/B |
| ALL#16 | *TCF3-PBX1*^b^ | 1A/B | **ALL#38** | *ETV6-RUNX1* | 4A/B |
| ALL#17 | *TCF3-PBX1*^b^ | 1B | **ALL#39** | *ETV6-RUNX1* | 4A/B |
| ALL#18 | *TCF3-PBX1* | 1A/B | **ALL#40** | B-other | 4A/B |
| ALL#19 | *TCF3-PBX1* | 1A/B | **ALL#41** | Hyperdiploid | 4A/B |
| ALL#20 | *TCF3-PBX1* | 1B | **ALL#42** | B-other | 4A/B |
| ALL#21 | *TCF3-PBX1* | 1B | **ALL#43** | *ETV6-RUNX1* | 4A/B |
| ALL#22 | *TCF3-PBX1* | 1A/B | **ALL#44** | B-other | 4A/B |

**Supplementary table 3. Ability of different passages MSCs to differentiate into adipo-, osteo-, and chondrocytes.** +++, ++, +, +/-, - indicate very efficient, efficient, well, intermediate, and poor differentiation of MSCs into adipo-, osteo-, chondrocytes, resp. p indicates MSC passage number.

**Adipocytes:**

| MSC# | Derived from | ALL subtype | p5 | p8 | p12 | p16 |
| --- | --- | --- | --- | --- | --- | --- |
| MSC#1 | Leukemic bone marrow, day 0 | B-other, B-ALL | **+++** | **+++** | **+++** | **++** |
| MSC#4 | Leukemic bone marrow, day 0 | B-other, B-ALL | **+++** | **+++** | **+++** | **++** |
| MSC#8 | Leukemic bone marrow, relapse | High hyperdiploid, B-ALL | **+++** | **+++** | **+++** | **++** |
| MSC#9 | Leukemic bone marrow, relapse | *MLL-AF4*, B-ALL | **+++** | **+++** | **++** | **++** |
| MSC#10 | Leukemic bone marrow, day 0 | T-ALL | **+++** | **+++** | **+++** | **++** |
| MSC#11 | Leukemic bone marrow, relapse | *ETV6-RUNX1*, B-ALL | **+++** | **+++** | **+++** | **++** |
| MSC#12 | Leukemic bone marrow, day 0 | *ETV6-RUNX1*, B-ALL | **++** | **++** | **++** | **++** |

**Osteocytes:**

| MSC# | Derived from | ALL subtype | p5 | p8 | p12 | p16 | p20 |
| --- | --- | --- | --- | --- | --- | --- | --- |
| MSC#1 | Leukemic bone marrow, day 0 | B-other, B-ALL | **+** | **+** | **+** | **-** | **-** |
| MSC#4 | Leukemic bone marrow, day 0 | B-other, B-ALL | **+++** | **+++** | **+++** | **++** | **++** |
| MSC#8 | Leukemic bone marrow, relapse | High hyperdiploid, B-ALL | **+++** | **+++** | **+++** | **++** | **-** |
| MSC#9 | Leukemic bone marrow, relapse | *MLL-AF4*, B-ALL | **+++** | **+++** | **++** | **++** | **-** |
| MSC#10 | Leukemic bone marrow, day 0 | T-ALL | **++** | **++** | **+/-** | **-** | **-** |
| MSC#11 | Leukemic bone marrow, relapse | *ETV6-RUNX1*, B-ALL | **+++** | **+++** | **+++** | **+** |  |
| MSC#12 | Leukemic bone marrow, day 0 | *ETV6-RUNX1*, B-ALL | **++** | **++** | **++** | **-** | **-** |

**Chondrocytes:**

| MSC# | Derived from | ALL subtype | p5 | p8 | p12 | p16 | p20 |
| --- | --- | --- | --- | --- | --- | --- | --- |
| MSC#1 | Leukemic bone marrow, day 0 | B-other, B-ALL | **+++** | **++** | **-** | **-** | **+** |
| MSC#4 | Leukemic bone marrow, day 0 | B-other, B-ALL | **+++** | **+++** | **+** | **+/-** | **-** |
| MSC#8 | Leukemic bone marrow, relapse | High hyperdiploid, B-ALL | **+** | **+** | **++** | **+++** | **++** |
| MSC#9 | Leukemic bone marrow, relapse | *MLL-AF4*, B-ALL | **-** | **++** | **+/-** | **+** | **-** |
| MSC#10 | Leukemic bone marrow, day 0 | T-ALL | **++** | **+++** | **++** | **+/-** | **-** |
| MSC#11 | Leukemic bone marrow, relapse | *ETV6-RUNX1*, B-ALL | **+++** | **+++** | **+** | **+** |  |
| MSC#12 | Leukemic bone marrow, day 0 | *ETV6-RUNX1*, B-ALL | **++** | **-** | **+/-** | **+** | **-** |

**Supplementary table 4. Survival benefit of BCP-ALL cells upon MSC co-culture.** ^a,b^ Two independent cultures with the same patient cells.

| ALL# | ALL subtype | Viability after thawing (percentage) | ALL mono (percentage CD19+) day 5 | ALL plus MSC mean (percentage CD19+; n= 4) day 5 | Benefit mean (percent point) | Benefit  standard dev | Benefit  qualitative |
| --- | --- | --- | --- | --- | --- | --- | --- |
| ALL#1 | *ETV6-RUNX1*^a^ | 87.5 | 44.4 | 61.5 | 17.2 | 2.7 | benefit |
| ALL#2 | *ETV6-RUNX1*^a^ | 94.2 | 58.4 | 66.3 | 7.9 | 5.6 | benefit |
| ALL#3 | *ETV6-RUNX1* | 95 | 64.5 | 75.0 | 10.4 | 1.6 | benefit |
| ALL#4 | *ETV6-RUNX1* | 97.6 | 26.5 | 37.0 | 10.5 | 1.9 | benefit |
| ALL#5 | *ETV6-RUNX1* | 91.4 | 61.7 | 78.2 | 16.6 | 3.5 | benefit |
| ALL#6 | *ETV6-RUNX1* | 95.1 | 36.1 | 39.5 | 3.5 | 1.7 | minimal benefit |
| ALL#7 | *ETV6-RUNX1* | unknown | 69.8 | 79.9 | 10.1 | 3.3 | benefit |
| ALL#8 | *ETV6-RUNX1* | 95.4 | 12.2 | 15.9 | 3.7 | 2.4 | minimal benefit |
| ALL#9 | *ETV6-RUNX1* | 91.8 | 42.3 | 48.3 | 6.0 | 0.8 | benefit |
| ALL#10 | *ETV6-RUNX1* | 93.9 | 17.7 | 30.2 | 12.5 | 2.0 | benefit |
| ALL#11 | Hyperdiploid | 91.9 | 0.4 | 0.8 | 0.4 | 0.2 | no benefit |
| ALL#12 | Hyperdiploid | 91 | 11.8 | 19.7 | 7.9 | 2.1 | benefit |
| ALL#13 | Hyperdiploid | 89.8 | 22.0 | 27.5 | 5.5 | 1.8 | benefit |
| ALL#14 | Hyperdiploid | 94 | 2.1 | 6.1 | 4.0 | 1.6 | minimal benefit |
| ALL#15 | *TCF3-PBX1* | 94.2 | 22.2 | 22.4 | 0.3 | 0.3 | no benefit |
| ALL#16 | *TCF3-PBX1*^b^ | 93.2 | 6.0 | 17.2 | 11.2 | 2.2 | benefit |
| ALL#17 | *TCF3-PBX1*^b^ | 93 | 0.5 | 6.0 | 5.5 | 2.4 | benefit |
| ALL#18 | *TCF3-PBX1* | 82 | 2.0 | 13.1 | 11.2 | 4.0 | benefit |
| ALL#19 | *TCF3-PBX1* | 81.9 | 2.2 | 5.9 | 3.7 | 1.3 | minimal benefit |
| ALL#20 | *TCF3-PBX1* | 83 | 2.0 | 9.5 | 7.5 | 1.6 | benefit |
| ALL#21 | *TCF3-PBX1* | unknown | 0.1 | 2.9 | 2.8 | 1.5 | minimal benefit |
| ALL#22 | *TCF3-PBX1* | 64.3 | 0.2 | 1.2 | 0.9 | 0.4 | no benefit |
| ALL#23 | B-other (*BCR-ABL1*-like) | 87.8 | 13.8 | 15.1 | 1.2 | 1.7 | no benefit |
| ALL#24 | B-other | 91 | 14.7 | 14.7 | 0.1 | 0.4 | no benefit |
| ALL#25 | B-other | 85 | 21.8 | 20.6 | -1.1 | 1.3 | no benefit |
| ALL#26 | *BCR-ABL1* | 84 | 0.8 | 25.2 | 24.4 | 11.0 | benefit |
| ALL#27 | *BCR-ABL1* | unknown | 0.4 | 8.6 | 8.2 | 0.7 | benefit |
| ALL#28 | *BCR-ABL1* | unknown | 1.8 | 12.2 | 10.4 | 2.6 | benefit |
| ALL#29 | *BCR-ABL1* | 90.1 | 1.0 | 3.0 | 2.0 | 0.5 | minimal benefit |
| ALL#30 | *BCR-ABL1* | 82.7 | 11.7 | 7.6 | -4.1 | 0.8 | worse |
| ALL#31 | *KMT2A-AF4* | 44 | 24.8 | 38.7 | 14.0 | 2.1 | benefit |
| ALL#32 | *KMT2A-AF4* | 65.4 | 7.6 | 28.6 | 21.0 | 1.9 | benefit |
